# Annotated bacterial chromosomes from frame-shift-corrected long read metagenomic data

**DOI:** 10.1101/511683

**Authors:** Krithika Arumugam, Caner Bağci, Irina Bessarab, Sina Beier, Benjamin Buchfink, Anna Gorska, Guanglei Qiu, Daniel H Huson, Rohan BH Williams

**Author notes:** Equal contributor (Daniel H Huson, Rohan BH Williams).

## Abstract

**Background:** Short-read sequencing technologies have long been the work-horse of microbiome analysis. Continuing technological advances are making the application of long-read sequencing to metagenomic samples increasingly feasible.

**Results:** We demonstrate that whole bacterial chromosomes can be obtained from a complex community, by application of MinION sequencing to a sample from an EBPR bio-reactor, producing 6Gb of sequence that assembles in to multiple closed bacterial chromosomes. We provide a simple pipeline for processing such data, which includes a new approach to correcting erroneous frame-shifts.

**Conclusions:** Advances in long read sequencing technology and corresponding algorithms will allow the routine extraction of whole chromosomes from environmental samples, providing a more detailed picture of individual members of a microbiome.

## Background

Second generation sequencing has been the work-horse of metagenomic analysis of microbiomes, with typical studies based on hundreds of millions of short reads [1, 2]. While the taxonomic and functional binning of short metagenomics read data are reasonably straight-forward computational problems [3], much recent work has focused on the challenge of assembling and binning metagenomic contigs, a procedure which provides invaluable working models of the genomes of member species [4]. However, the assembly of whole bacterial chromosomes from short metagenomic reads has proven to be an all but impossible task.

Third generation sequencing promises to allow the extraction of whole genomes from environmental samples with ease [5]. This promise is now beginning to be fulfilled. Here we report on the results of a single ONT MinION run on a microbial community from an enrichment bioreactor targeting polyphosphate accumulating organisms (PAO), that had been inoculated with activated sludge from a full-scale water reclamation plant in Singapore.

## Results

Running a MinION sequencer for one day, we obtained 695, 000 long reads with an average length of 9kb, totaling approximately 6Gb of sequence (Table S1). Using Unicycler [6], we assembled these into 1, 702 contigs (LR-contigs) of average length 61kb (Table S2). We observed 10 contigs over 1 Mb in length, including five circular contigs between 2.7 and 4.2 Mb long (see Figure 1A). In principle, long read assembly procedures could generate complete genomes *de novo*, without the need for complex contig binning procedures, and accordingly we designed tools and analyses to determine the extent to which such long contigs represented genomes of member species of the community. Our analyses are based on 1) analysis of genome completeness and quality; 2) whole genome comparison to reference genomes, and 3) comparison with metagenome-assembled genomes recovered from short read sequence generated from the same DNA sample.

**Figure 1.**
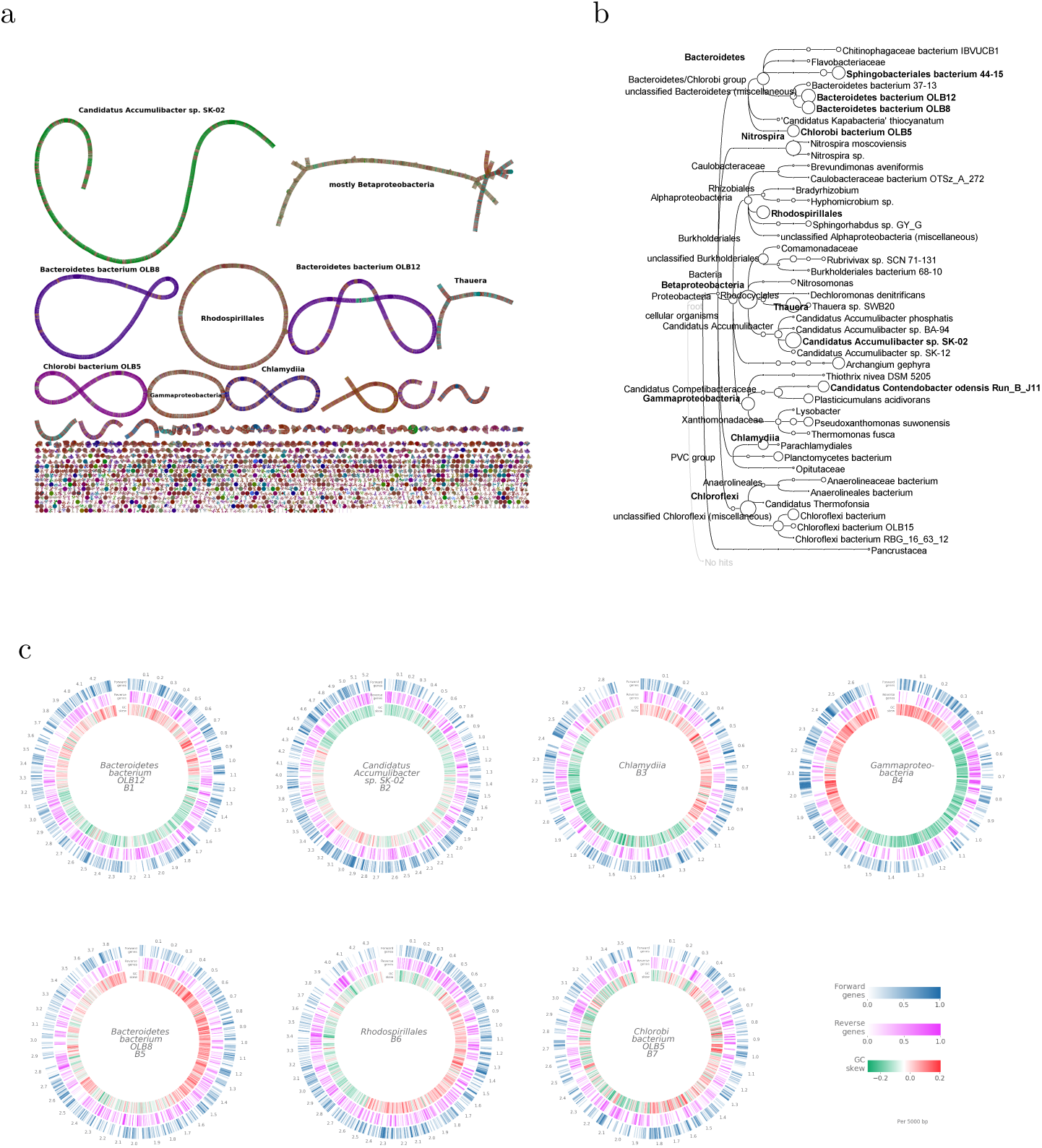
Summary of results. a: Bandage [19] visualization of the Unicycler assembly graph before final segmentation into contigs. The largest connected components are labeled by the corresponding taxonomic bins and the nodes are colored by the MEGAN taxonomic classification of the long reads. The seven longest linear and circular components correspond to the seven LR-chromosomes. b: MEGAN-LR taxonomic binning: nodes are scaled to indicate the number of aligned bases in each bin. Bins that are more than 50% complete are shown in bold. c: Annotation of the seven LR-chromosomes, labeled by the corresponding taxonomic bins. The three circular tracks indicate the genes annotated by Prokka on the forward strand (blue) and reverse strand (pink), and GC-skew (green and red indicate lower or higher than average GC content, respectively).

Long reads, and, to a slightly lesser degree, LR-contigs, suffer from a high rate of erroneous insertions and deletions, which lead to frame-shifts in translated alignments (in the present case, we observed 7-15 frame-shifts per kb in coding sequence). For this reason, genome evaluation tools (such as CheckM [7]) and annotation workflows (such as Prokka [8]), which typically employ translated alignments, perform poorly on current long read data.

To address this deficiency, we have developed a two-step frame-shift correction technique. First, we have modified DIAMOND [9] (v 0.9.23) to perform a *frame-shift aware* DNA-to-protein alignment [10] of the sequences against the NCBI-nr protein reference base [11]. Second, based on the location of frame-shifts reported in the alignments, we insert N’s into the sequences so as to maintain the frame (see Figure 2B). Thus sequences corrected in this way can be evaluated and annotated using conventional genome quality and annotation tools.

**Figure 2.**
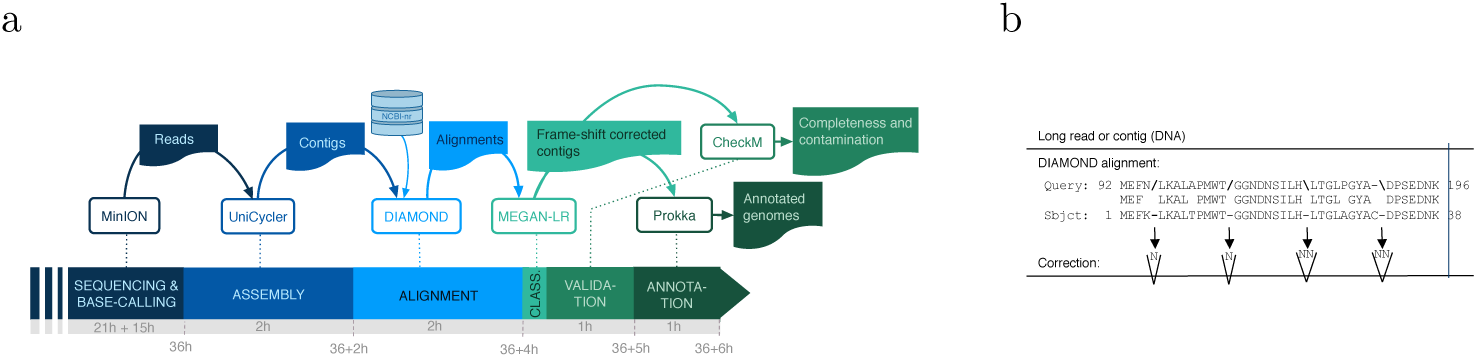
Analysis. a: Long read analysis pipeline shown from left to right. MinION sequencing produces a set of reads. These are assembled into contigs using Unicycler and aligned against the NCBI-nr database using DIAMOND. The contigs and alignments are processed by MEGAN to perform taxonomic binning and also to produce frame-shift-corrected contigs. These are analyzed using CheckM and annotated using Prokka. The duration of each step is shown in wall-clock hours. MEGAN analysis took less than 10 minutes. b: Frame-shift correction: in frame-shift alignments, forward slashes and backward slashes indicate a frame decrease, or increase, by one, respectively. Correction is performed by inserting one or two unspecified nucleotides into the sequence, respectively.

We performed initial taxonomic analysis of all LR-contigs using MEGAN-LR [12] (v 6.13.3), obtaining 106 taxonomic bins at different taxonomic ranks (see Figure 1B and Table S3). To determine whether these taxonomic bins might harbor complete genomes, we applied CheckM to the set of frame-shift corrected LR-contigs contained in each taxonomic bin. This analysis indicates that 14 of the bins are more than 50% complete. Of these, six fulfill the definition of a “high quality draft” metagenome-assembled genome (namely, completeness *>* 90% and contamination *<* 5%). For purposes of this paper, we also consider the seventh bin listed in Table 1 as high quality, as it consists of only one circular LR-contig and is of chromosomal length. There are four additional bins that reach the level of “medium quality draft” (completeness *>* 50% and contamination *<* 10%)[13].

**Table 1.** Summary of results. For all 14 taxonomic bins B1–B14 that CheckM deems *≥* 50% complete (a), and -in cases where the bin contains more than one contig-also for the longest contig, in descending order of assembly quality, we report: (b) the number of contigs produced by Unicycler, (c) the total number of bases, (d) the number of bases aligned by DIAMOND to some protein reference, (e) the average coverage by long reads (based on the longest contig), (f) the %-completeness and (g) %-contamination reported by CheckM, and (h)–(j), the number of rRNA, tRNA and coding sequences reported by Prokka, respectively.

|  | (a)<br>DIAMOND+MEGAN<br>taxonomic bin | (b)<br>Unicycler<br>contigs | (c)<br>Total<br>(Mb) | (d)<br>Aligned<br>(Mb) | (e)<br>Average<br>coverage | (f)<br>CheckM<br>Complete. | (g)<br>Contam. | (h)<br>rRNA | (i)<br>Prokka<br>tRNA | (j)<br>CDS |
| --- | --- | --- | --- | --- | --- | --- | --- | --- | --- | --- |
| High quality draft genomes: |  |  |  |  |  |  |  |  |  |  |
| B1 | *Bacteroidetes bacterium* OLB12 | 1 | 4.2 | 3.5 | 57.3 | 95% | 0.1% | 6 | 39 | 4,163 |
| B2 | *Candidatus Accumulibacter* SK-02 | 1 | 5.2 | 4.1 | 384.2 | 94% | 0.6% | 4 | 53 | 4,915 |
| B3 | *Chlamydia* (class) | 1 | 2.8 | 1.8 | 48.8 | 94% | 2% | 6 | 39 | 3,387 |
| B4 | *Gammaproteobacteria* (class) | 43 | 4.7 | 3.0 |  | 93% | 2% | 6 | 52 | 4,833 |
|  | -longest contig |  | 2.7 | 1.6 | 25.1 | 93% | 0.2% | 3 | 40 | 3,359 |
| B5 | *Bacteroidetes bacterium* OLB8 | 1 | 3.8 | 3.0 | 52.1 | 93% | 1% | 6 | 37 | 3,394 |
| B6 | *Rhodospirillales* (order) | 1 | 4.4 | 3.0 | 29.5 | 92% | 0.5% | 3 | 47 | 4,015 |
| B7 | *Chlorobi bacterium* OLB5 | 1 | 3.5 | 2.5 | 38.7 | 88% | 1% | 3 | 41 | 4,131 |
| Medium quality draft genomes: |  |  |  |  |  |  |  |  |  |  |
| B8 | *Thauera* (genus) | 25 | 4.6 | 4.0 |  | 89% | 4% | 12 | 64 | 4,040 |
|  | -longest contig |  | 0.8 | 0.7 | 32.7 | 14% | 0% | 0 | 5 | 672 |
| B9 | *Sphingobacteriales bacterium* 44-15 | 59 | 3.2 | 2.8 |  | 76% | 1% | 2 | 17 | 2,953 |
|  | -longest contig |  | 0.2 | 0.1 | 10.2 | 0% | 0% | 0 | 0 | 172 |
| B10 | *Bacteroidetes* (phylum) | 43 | 3.9 | 2.6 |  | 72% | 7% | 1 | 12 | 1,997 |
|  | -longest contig |  | 1.2 | 0.8 | 14.1 | 32% | 0% | 0 | 3 | 807 |
| B11 | *Candidatus Contendobacter* B J11 | 39 | 2.5 | 2.0 |  | 59% | 9% | 2 | 37 | 2,668 |
|  | -longest contig |  | 0.3 | 0.3 | 15.4 | 19% | 0% | 0 | 7 | 295 |
| Low quality draft genomes: |  |  |  |  |  |  |  |  |  |  |
| B12 | *Betaproteobacteria* (class) | 111 | 6.6 | 5.5 |  | 89% | 79% | 6 | 71 | 4,655 |
|  | -longest contig |  | 0.4 | 0.3 | 37.1 | 10% | 0% | 0 | 1 | 372 |
| B13 | *Nitrospira* (genus) | 34 | 4.2 | 3.7 |  | 83% | 13% | 0 | 6 | 563 |
|  | -longest contig |  | 1.1 | 0.9 | 17.6 | 27% | 0% | 0 | 2 | 99 |
| B14 | *Chloroflexi* (phylum) | 151 | 5.4 | 4.3 |  | 71% | 29% | 0 | 11 | 3,565 |
|  | -longest contig |  | 0.2 | 0.2 | 13.3 | 8% | 0% | 0 | 1 | 86 |

In all seven high-quality bins, the CheckM results derive from a single long contig, of length 2.7 – 5.2Mb, with the numbers of cognate rRNA and tRNA genes, and protein coding genes, as reported by Prokka, all lying within the range usually seen for bacterial genomes (see Table 1 and Table S3). Throughout this paper, we will refer to these long contigs as the seven *LR-chromosomes*.

From the seven high quality taxonomic bins, we obtained a near-complete LR-chromosome (number B2 in Table 1) that is binned to *Candidatus Accumulibacter*, a phosphate accumulating organism (PAO) that operates in waste-water treatment plants and is the target of our enrichment protocol [14]. Two circular LR-chromosomes (B1 and B5) are binned to the species *Bacteroidetes bacterium* OLB8 and OLB12, both of which were originally recovered as metagenome-assembled genomes from a partial-nitritation anammox (PNA) bioreactor community, where they are are thought to function as aerobic heterotrophs [15]. All three of these LR-chromosomes align end-to-end to their corresponding reference genomes (see Figure 3).

**Figure 3.**
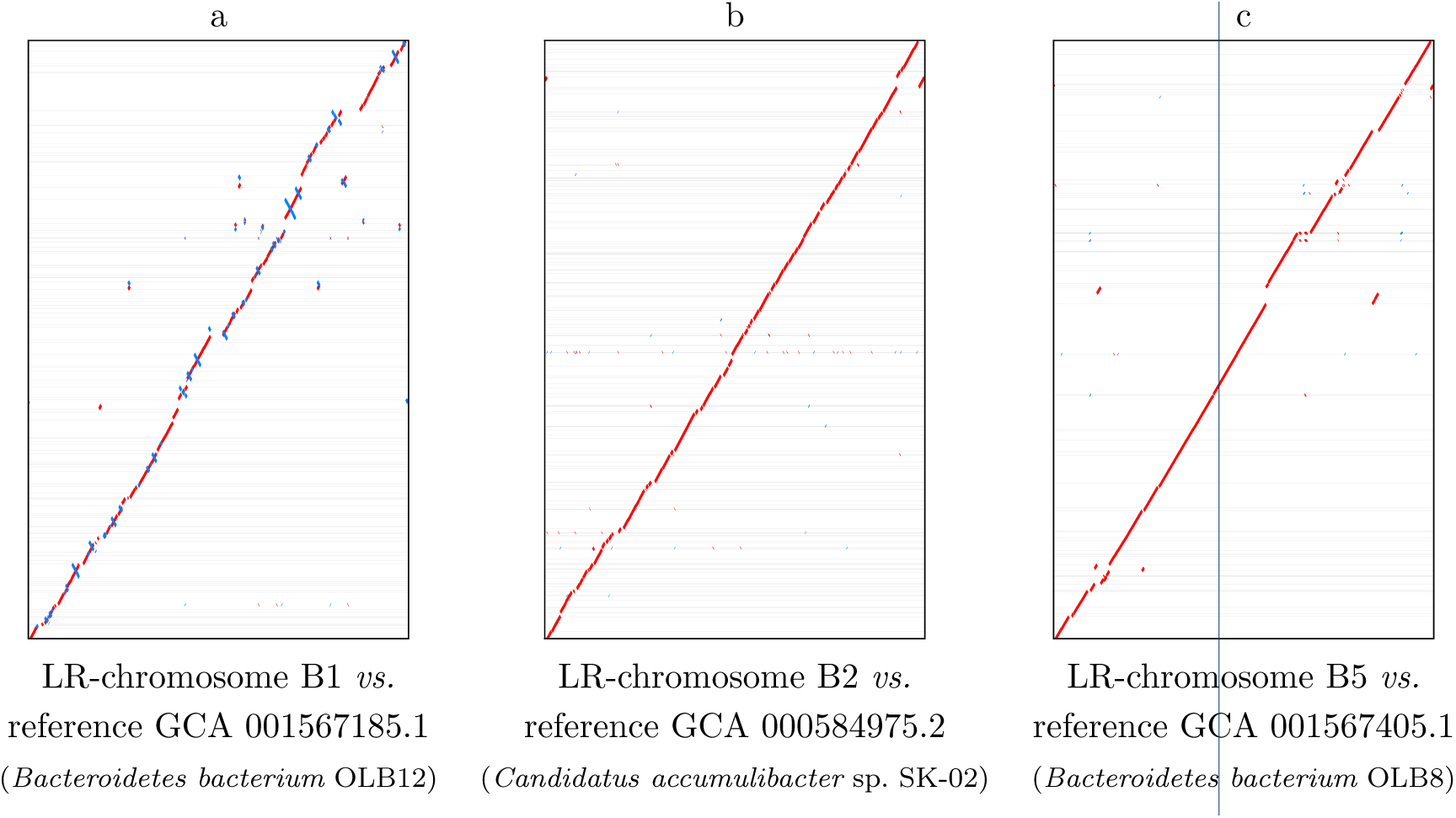
Dot-plots for the three LR-chromosomes that have high similarity to reference genome assemblies, (a) B1, (b) B2 and (c) B5. The LR-chromosomes are represented by the *x*-axis and the corresponding reference assembly is represented by the *y*-axis. Forward alignments are shown in red, whereas reverse complemented alignments are shown in blue, gray lines indicate contig boundaries in the reference assemblies.

The remaining four are closed circular chromosomes that do not align to any current reference genome and thus most likely represent novel organisms. One of these (B3) that is binned to the class of *Chlamydiia*. Although normally considered an obligate intracellular pathogen in humans, members of the phylum *Chlamydiae* are known to occur in microeukaryotes that occur as predators in such reactor communities [16]. Another (B6) is binned to *Rhodospirillales* and contains a 16S sequence that maps to the genus *Defluviicoccus*. Some members of this genus compete with PAO for carbon sources and are commonly observed in PAO enrichment reactors [17]. Another LR-chromosome (B4) is binned to class *Gammaproteobacteria*. Finally, we obtained an LR-chromosome (B7) that is binned to *Chlorobi bacterium OLB5*, an organism previously observed in waste-water [15].

For all seven LR-chromosomes, Silva analysis [18] of the contained 16S sequences confirm the taxon bin assignment obtained by MEGAN analysis.

Solely for the purpose of verification, we also produced a second independent set of paired reads from the same DNA aliquot using Illumina short read sequencing. First, we used the short-read clone coverage to detect potential break-points in the assemblies of seven LR-chromosomes that might indicate long read assembly errors, and found 11. All but one of these positions have very good long-read coverage, making an assembly error unlikely at these positions. Second, we assembled the short reads and aligned the short-read contigs against the long-read contigs, and this comparison shows a very high degree of co-linearity (see Figure 5). Third, we performed metagenomic binning of the short-read contigs and compared the short-read bins with the long-read chromosomes, obtaining a high degree concordance between the two assemblies (see Figures 6 and 7).

**Figure 6.**
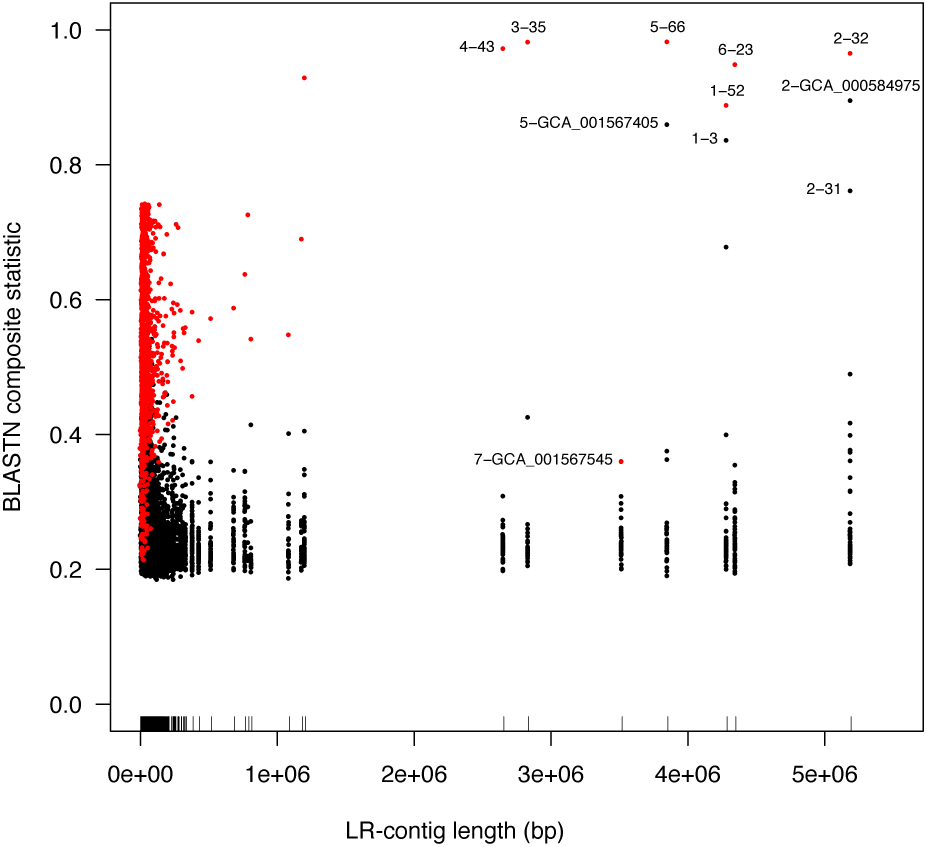
Overview of alignment statistics between LR-contigs, on the one hand, and short read assembly bins (SR bins) and reference genomes, on the other. The *x*-axis shows the length of each LR-contig with the position of each tagged with a tick on the axis; the *y*-axis show the value of the BLASTN summary statistic, *sstat* (see Methods) and data points represent pairs of LR-chromosomes and SR-bins (or references genomes). Selected pairs of tags with high values of *sstat* are highlighted with “ ⟨LR-chromosome.id ⟩-⟨SR-bin.id ⟩” for comparisons to short read assembly bins or “ ⟨LR-chromosome.id ⟩ - ⟨GCA id ⟩” for comparisons to references. Within each set of the seven LR-chromosome alignments, the pair with the maximum *sstat* value is shown in red. All LR-chromosomes have highly aligned counterpart SR-bins, with the exception of LR-chromosome 5. Further data on individual LR-chromosomes alignments are shown in Fig 7.

**Figure 7.**
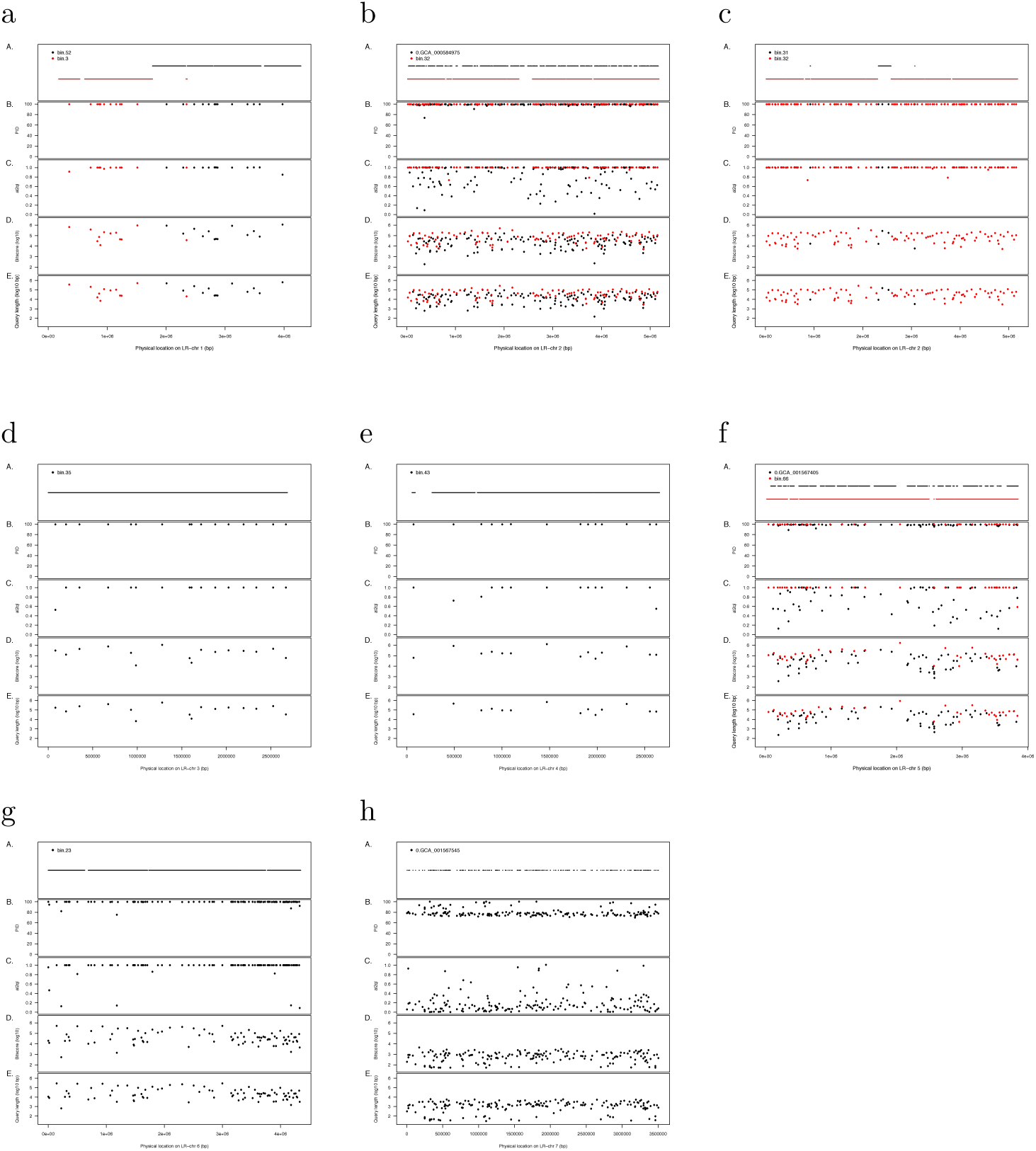
Alignment statistics for SR-contigs against LR-chromosomes. The LR-chromosome is represented by the *x*-axis. Each plot contains five panels, from top to bottom, representing: (A) the locations of alignments to the LR-chromosome, (B) the corresponding percent identity, (C) alignment-length to query-length ratio, (D) alignment length and (E) query length. The colors red and black are used to distinguish between alignments to different SR-bins or reference genomes, as described in the text.

**Figure 5.**
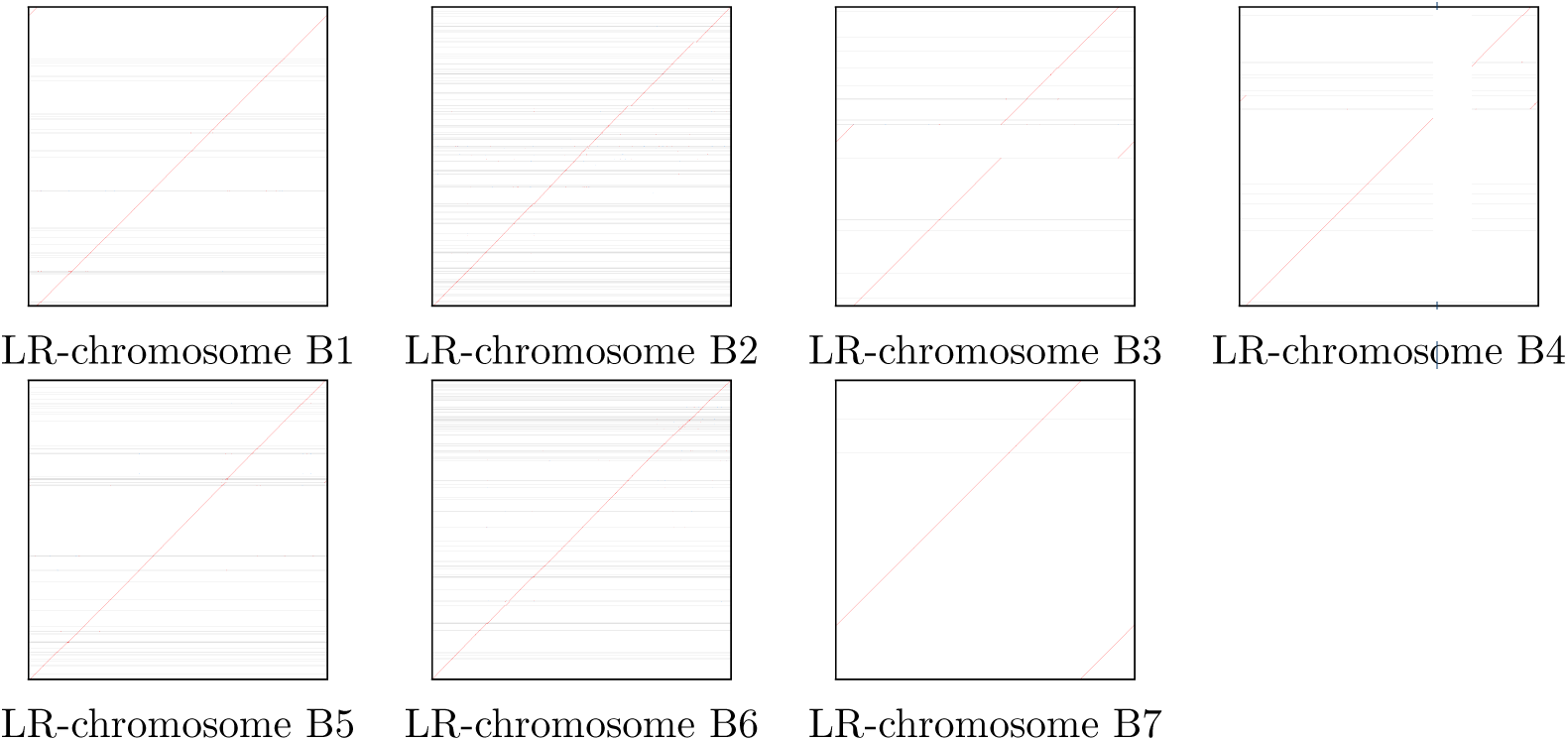
For each of the seven LR-chromosomes (B1–B7), we show a dot plot comparison of the chromosome (*x*-axis) with the set of all SR-contigs (*y*-axis).

## Discussion

In this study, a single run of a Nanopore MinION device on a complex bioreactor community gave rise to a high coverage (384x) of the target polyphosphate accumulating organism, *Candidatus Accumulibacter*, but also 10-60x coverage for 13 other taxa. From this data, in total, seven high quality draft genomes were obtained, six of which as closed circular chromosomes. Only three of these draft genomes have corresponding reference genomes at NCBI. In all three cases, the LR-chromosomes display a major improvement in continuity over the reference genomes, which were obtained by metagenomic assembly of short reads.

A potential concern might be that the reported megabase-sized contigs might be chimeric or otherwise incorrect. The results reported by CheckM and Prokka suggest that these sequences are entirely consistent with being complete bacterial chromosomes. Moreover, our comparison with a set of short reads sequenced from the same DNA provides further evidence that the reported LR-chromosomes are correct, and that an extremely high degree of recapitulation is obtained when compared to draft genomes obtained from the same DNA extraction.

One current issue with long read sequencing technologies is that they produce a significant rate of erroneous insertions and deletions, which cause problems when performing translated alignments. Our work suggests that frame-shift aware alignment techniques can be used to reduce such problems.

## Conclusions

This work suggests that it is now possible to obtain complete bacterial chromosomes from a complex microbial community using Nanopore sequencing. We provide a straight-forward pipeline for processing such data. It performs assembly, alignment against NCBI-nr, taxonomic binning, frame-shift correction, bin quality analysis and annotation, in less than six hours (see Figure 2A).

The application of long read sequencing techniques promises to allow the routine extraction of whole chromosomes from environmental samples, providing a much more detailed picture of individual members of a microbiome.

## Methods

### EPBR bioreactor

A sequencing batch reactor (SBR) with 5.4 L working volume was inoculated with activated sludge from an EBPR mother reactor. A slow feeding strategy was applied for the reactor operation, which has been shown to benefit the proliferation of *Ca.* Accumulibacter [20]. The SBR was operated in six hour cycles, including 60 min feeding, 20 min anaerobic, 180 min aerobic, and a 100 min settling/decant stage. In each cycle, 2.35 L of synthetic waste-water composed of 0.53 L of solution A (containing 1.02g/L NH_4_Cl, 1.2g/L MgSO_4_ 7H_2_O, 0.01g/L peptone, 0.01g/L yeast extract and 6.8 g/L sodium acetate) and 1.82 L of solution B (0. 312g/L K_2_HPO_4_ 3H_2_O, 0.185 g/L KH_2_PO_4_, 0.75 mg/L FeCl_3_ 6H_2_O 0.015 mg/L CuSO_4_ 5H_2_O, 0.03 mg/L MnCl_2_, 0.06 mg/L ZnSO_4_, 0.075 mg/L CoCl_2_, 0.075 mg/L H_3_BO_3_, 0.09mg/L KI and 0.06 mg/L Na_2_MoO_4_ 2H_2_O) (modified from [21]) was introduced into the reactor. The reactor was operated at 30 °C with an hydraulic retention time (HRT) and a solid retention time (SRT) of 12 h and 11 days, respectively. The pH was controlled at 7.00–7.60 with DO levels maintained at 0.8-1.2 mg/L during the aerobic phase. The SBR achieved P-release of 180–200 mg/L with complete P removal observed after six month operation. The reactor was sampled on day 267 of operation.

### DNA extraction

Genomic DNA was extracted from the sampled biomass with the FastDNA™SPIN kit (MP Biomedicals) for Soil, using 2*×* bead beating with a FastPrep homogenizer (MP Biomedicals). The DNA was then size-selected on a Blue Pippin DNA size selection device (SageScience) using a BLF-7510 cassette with high pass filtering with a 8 kb cut off.

### Nanopore sequencing

The sequencing library was constructed from approximately 4*µ*g of genomic DNA using the SQK-LSK 108 Ligation Sequencing Kit (Oxford Nanopore Technologies). Sequencing was performed on a MinION Mk1B instrument (Oxford Nanopore Technologies) using a SpotON FLO MIN106 flowcell (FAH85393) and R 9.4 chemistry, running for approximately 24 hours. Data acquisition was performed using MinKNOW version 1.14.1 running on a HP ProDesk 600G2 computer (64–bit, 16Gb RAM, 2Tb SSD HD) running Windows10. Base-calling was performed using Albacore version 2.3.1. Adaptor trimming was performed using Porechop [22] with default settings. This produced 694, 955 reads of average length 9kb (range: 2bp– 66kb). A summary of the long read statistics is given in Supplementary Table S1.

### Long read assembly

Long read assembly was performed using Unicycler (v 0.4.6) running with default settings. Assembly of the 694, 955 long reads produced 1, 702 LR-contigs of average length 61kb (1.3kb - 5.2Mb). This took about two hours on a server. (All timings in this paper measured on a server with AMD Opteron(TM) Processor 6274, 64 x 2.2GHz, 512GB memory). A summary of the long read contig statistics is given in Supplementary Table S2.

### DIAMOND options for long reads

This paper introduces two new features in DIAMOND for use with error-prone long reads or contigs. First, the program now provides a *frame-shift mode* that performs frame-shift alignment of DNA sequences against a protein reference database [10]. This feature is activated using the command line option -F 15, which also sets the frame-shift dynamic programming penalty to a specific value, in this case 15.

Second, the program now provides the option to perform *range-culling*. This feature determines which alignments are reported to output. Without range-culling, the program reports the most significant alignments for the query, up to a given count or score, independent of their position along the query. With range-culling, the decision whether to report an alignment is made locally. By default, any alignment *A* found is reported, unless there exists another alignment *B* that covers at least 50% of *A* on the query and whose bit-score is significantly larger, by defaulting requiring that the score of *A* is less than 90% of the score of *B*. This feature is activated using the command line options `--range-culling` and `--top 10`.

### DIAMOND alignment

In preparation of running DIAMOND on the Unicycler LR-contigs, we downloaded the NCBI-nr database in November 2018, obtaining 177.6 million protein reference sequences. DIAMOND required about one hour to initially process the database.

DIAMOND was run on the LR-contigs with the following options: `--range-culling --top 10 -F 15 --outfmt 100 -c1 -b12 -t /dev/shm`. The program required 104 minutes to align the 1,703 LR-contigs against the NBCI-nr database and obtain 1.8 million alignments for 1,695 contigs.

### Frame-shift correction

In Figure 2B we illustrate how to correct frame-shift errors in a given query DNA sequence, based on an alignment computed by DIAMOND in frame-shift mode. In a frame-shift alignment, a ‘/’ in the alignment transcript indicates that the aligner decreased the current frame of the query sequence by 1 at the given position, whereas a ‘*\*’ indicates the current frame was decreased by 1, as in http://last.cbrc.jp/doc/lastal.html. To perform frame-shift correction, in the former case, we insert a single unspecified nucleotide ‘N’ into the query sequence, whereas in the latter case, we insert two unspecified nucleotides ‘NN’.

To perform this correction on a long read or LR-contig, we greedily select a maximal set of non-overlapping alignments for the whole query and use this set for correction. This is implemented in MEGAN.

### MEGAN analysis and frame-shift correction

The output file of DIAMOND was prepared for analysis with MEGAN using the program daa-meganizer, which is part of the MEGAN Community Edition suite, version 6.13.1. The following command line options were used:

~~~
--longReads --lcaAlgorithm longReads --lcaCoveragePercent
51 --readAssignmentMode alignedBases --acc2taxa
prot acc2tax-Nov2018X1.abin
~~~

The first three options select MEGAN’s long-read analysis mode and sets the amount of aligned sequence to be covered by a taxon during the LCA analysis to 51% [12]. The fourth option requests that the primary count associated with each taxon is the number of aligned reads contained in the contigs binned to that taxon. The final option instructs the program to use the November-2018 mapping of NCBI accessions to NCBI taxa. This “meganization” step took less than five minutes.

A summary of the taxon bins obtained by MEGAN analysis is given in Supplementary Table S3.

Frame-shift correction was performed on all LR-contigs using MEGAN’s `Export Frame-Shift Corrected Reads…` menu item and the resulting sequences were saved into taxon-specific files, in just over two minutes.

### CheckM

The frame-shift corrected bins were analyzed for their completeness and contamination using CheckM (v1.0.12) in `lineage wf mode`. Data files for CheckM were downloaded on 26.11.2018 from https://data.ace.uq.edu.au/public/CheckM_ databases. The full output of CheckM is provided in Supplementary Table S4.

### Prokka

We annotated the frame-shift-corrected bins using Prokka (v1.12) in metagenome annotation mode without specifying taxa. The taxonomic database for this version of Prokka is based on Rfam 1.12.

### 16S analysis

For all seven LR-chromosomes, we extracted all 16S sequences annotated by Prokka and performed taxonomic classification of them using Silva [18], obtaining the correspondence between the MEGAN assignments and the Silva assignments (note that Ignavibacteriaceae appears within the Chlorobi group in the NCBI taxonomy) reported in Table 2.

**Table 2.**
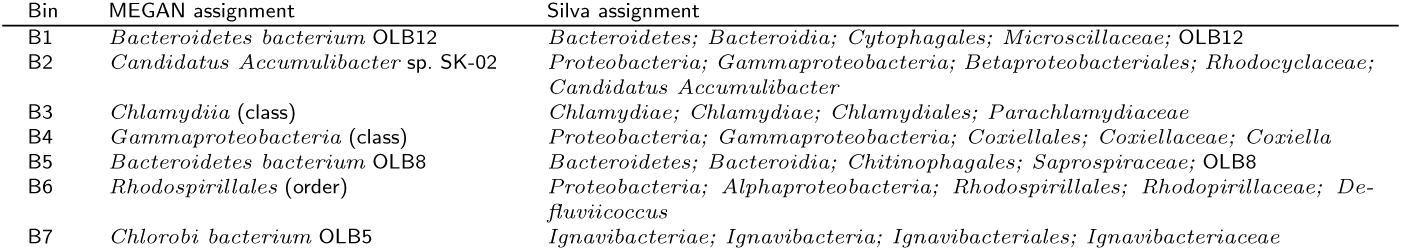
For all seven LR-chromosomes, we list the MEGAN and Silva taxonomic assignments.

All assignments were obtained using a threshold of 95% identity, except for the case of bin B4, where a lower threshold of 90% identity was needed to obtain an assignment.

### Comparison with genomic references

For each of the seven LR-chromosomes, we determined the reference taxon that occurs the most times in DIAMOND alignments of the contig against NCBI-nr. We then aligned the LR-chromosomes to the corresponding reference assemblies using Minimap2 (v2.14-r883) with parameters `-cx asm20 -t32 --secondary=yes -P`. We found a significant level of DNA similarity in three cases, which we summarize here as dot plots (see Figure 3). The other four LR-chromosomes did not align to their corresponding reference sequences (less than 1% of the total chromosome covered by an alignment), or, indeed, to any genome in the whole of NCBI-nt.

### Repeat analysis

We used Minimap2[23] to align all seven LR-chromosomes against themselves with parameters `-cx asm10 -t32 --secondary=yes -P` to find repeated regions in them. The option -c generates CIGAR strings in the output, `-x asm10` is a preset of parameters for comparing assemblies with up to 10% divergence, `-t32` sets the number of threads, `--secondary=yes` reports secondary alignments (by default Minimap2 reports only the best alignment), and `-P` retains all chains and attempts to elongate them. We then marked the positions that are within alignments of length equal to or greater than 500 in a contig to itself as repeat regions.

In order to check whether the repeat rates obtained for our contigs are typical for bacterial genomes, we performed the same analysis on all complete bacterial genomes in RefSeq (downloaded on 01.06.2018). Figure 4 suggests that the seven LR-chromosomes have repeat-rates that are similar to those observed for complete bacterial genomes in RefSeq.

**Figure 4.**
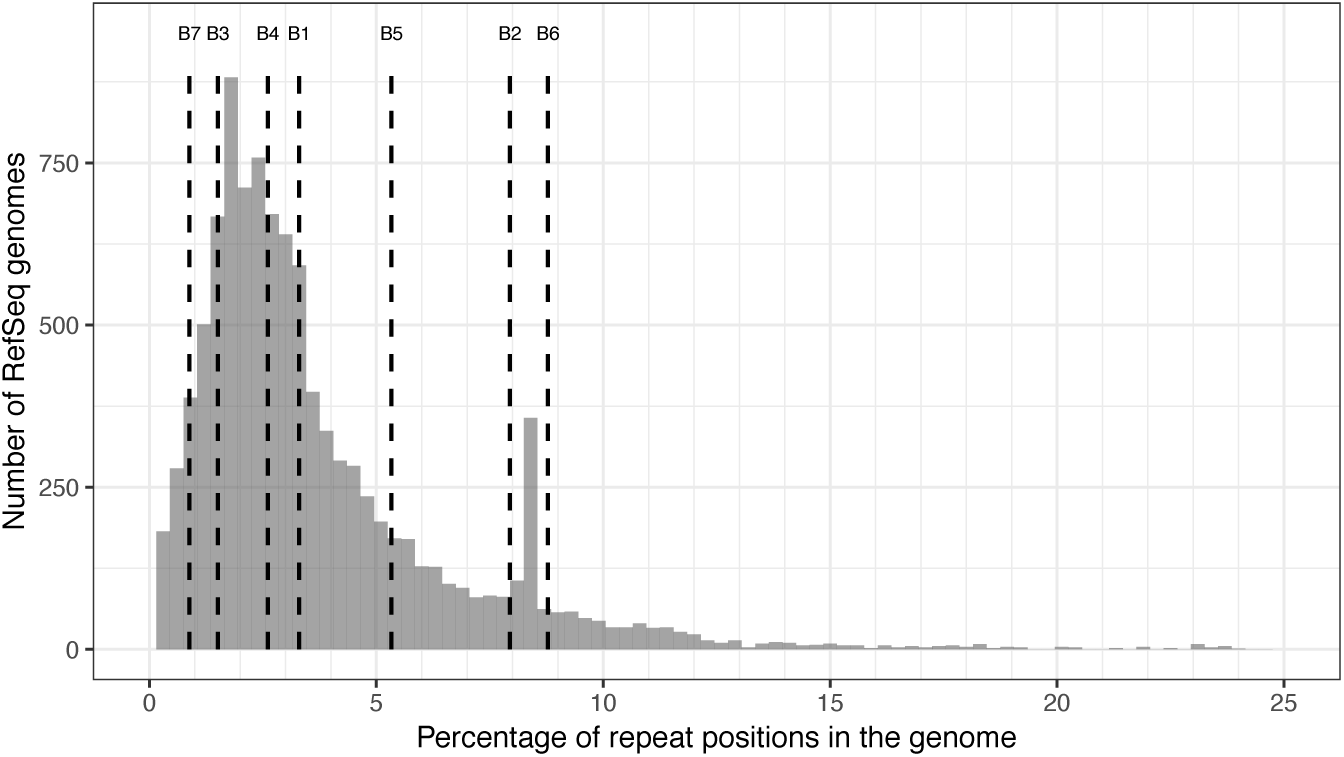
Distribution of repeat rates in all complete bacterial genomes in RefSeq. Vertical lines show the repeat rate of the seven LR-chromosomes. Additional 17 data points that have a repeat percentage above 25% are not shown.

### Additional short read sequencing

To support the evaluation of the long read contigs, we performed additional short read sequencing from the same sample. Genomic DNA Library preparation was performed using a modified version of the Illumina TruSeq DNA Sample Preparation protocol and sequenced on an Illumina HiSeq 2500 sequencing using a read length of 251 bp (paired-end). The raw gDNA FASTQ files were processed using cutadapt (v 1.14) with Python 3.6.3 in paired end mode (with default arguments except `-overlap 10 -m 30 -q 20,20`). We obtained 43,856,872 short reads in total. A summary of the short reads is provided in Supplementary Table S5.

### Break-point and coverage analysis using short reads

We aligned all short reads against the LR-contigs using Minimap2 [23] (options: `-2 -f 0 -t 32 -F 10000 -ax sr --secondary=yes -N 10000`). Then, considering each pair of reads a valid clone, if the two aligned reads have the correct orientation with respect to each other and a distance below 800, we determined the clone coverage of each LR-contig. Any stretch of LR-contig, for which the clone-coverage is zero, is considered a potential break-point. We identified 11. All but one of these are covered by multiple long reads, and so we assume that they are not indicative of a long read assembly error. The coordinates of the potential break-points are reported in Supplementary Table S6.

A comparison of the SR-coverage and LR-coverage of the 14 longest LR-contigs reported in Table 1 yields a strong positive correlation (Pearson R=0.9988), see Table 3.

**Table 3.** Comparison of LR- and SR-coverage. For each of the longest contigs in the 14 bins reported in Table 1, we report the average long-read coverage and average short-read coverage.

| Bin | LR-coverage | SR-coverage |
| --- | --- | --- |
| B1 | 57.3 | 117.5 |
| B2 | 384.2 | 707.8 |
| B3 | 48.8 | 107.6 |
| B4 | 25.1 | 56.2 |
| B5 | 52.1 | 109.9 |
| B6 | 29.5 | 56.1 |
| B7 | 38.7 | 90.2 |
| B8 | 32.7 | 60.5 |
| B9 | 10.2 | 23.2 |
| B10 | 14.1 | 27.7 |
| B11 | 15.4 | 22.3 |
| B12 | 37.1 | 66.0 |
| B13 | 17.6 | 28.7 |
| B14 | 13.3 | 18.9 |

### Assembly of short reads

The 43.86 million short reads were assembled using SPAdes-3.12.0 [24] (default parameters except `-meta -k 21,33,55,77,99,127 -t 30`). We obtained a total of 539,404 short-read contigs (SR-contigs) of at least 500bp in length. A summary of the SRC-contigs is provided in Supplementary Table S7.

### Comparison of SR-contigs and LR-chromosomes

To verify the correctness of the seven LR-chromosomes, we aligned them against the set of SR-contigs using Minimap2, as described in Repeat analysis section, and present the results as a dot-plot in Figure 5. These plots indicate a perfect concordance between the LR-chromosomes and corresponding SR-contigs. (What appear to be breaks in four of the diagonals are artifacts due to “wraparound” in the circular chromosomes.)

For each of the seven LR-chromosomes, we aligned all corresponding SR-contigs against the corresponding reference genomes (as described above) using Minimap2 and find significant alignments only for SR-contigs corresponding to the LR-chromosomes 1, 2 and 5. This supports the conclusion that only three of the LR-chromosomes are present in current reference databases.

### Metagenomic binning of short read assembly

Genome binning was performed using Metabat [25] (v2.12.1, using default parameters) on all SR-contigs that were at least 2kb in length, with bin evaluation being performed with CheckM (v1.0.11) (default parameters except `lineage wf -t 29`). This gave rise to 80 bins, of which 21 (26%) fulfill the definition of “high quality” and 14 (18%) are considered “medium quality” [13]. We screened for 16S genes within the SR-contigs using the USEARCH [26] module `--search16s` (v 10.0.240, 64 bit), and annotated these sequences using Silva.

### Comparison of SR-bins and LR-chromosomes

We used BLASTN [27] (version 2.4.0+) to examine the degree of sequence alignment between LR-contigs and SR-contigs sequences. We treated the LR-contigs as the subject sequences and the SR-contigs as the query sequences, using default parameters, and retained the best hit from each SR-contig to LR-contig alignment, if observed. Alignment statistics were then categorized by LR-contig and SR-bin memberships, and we then assessed the extent to which SR-contigs aligned to LR-contigs sequence, we utilized the following statistics 1) ratio of the alignment length to the length of the query (SR-contig) sequence (*al*2*ql*); 2) BLASTN identity (expressed as a proportion, not a percentage); 3) the proportion of the LR-contig that is covered by aligned SR-contigs and 4) the proportion of the SR-contigs in the bin that are aligned on the LR-contig. For each of these, we calculated their mean value across all alignments observed in each pairwise combination of LR-contig and SR-bin. As each of the four statistics just described is actually or effectively bounded between 0 and 1, if we calculate their overall mean (referred to hereafter as *sstat*), and select for values of *sstat* close to 1, in effect, selecting for SR-bins in which the majority of member contigs should tile a given LR-contig with a high degree of uniform alignment.

### Detailed comparison of SR-bins and LR-chromosomes

Here we provide a brief summary of the taxonomic annotation of LR-chromosome sequences and further highlight inter-relationships between LR-chromosomes, on the one hand, and SR-bins and reference genomes, on the other.

LR-chromosome 1 is contained within the MEGAN-LR taxonomic bin annotated to *Bacteroidetes bacterium OLB12*. This LR-chromosome is tiled by contigs from SR-bin 52 (medium quality “metagenome assembled genome” (MAG), with *sstat* = 0.88) and SR-bin 3 (*sstat* = 0.84), which cover the first third and second two-thirds of this LR sequence, respectively. SR-bin 52 is annotated by CheckM to UID2570, which is selective for members of phyla Chlorobi, Bacteroidetes and Ignavibacteriae. The precise taxonomic placement is not clear. See Figure 7a.

LR-chromosome 2 is contained within the MEGAN-LR taxonomic bin annotated to *Candidatus Accumulibacter sp. SK-02*, and is tiled (*sstat* = 0.97) by contigs in SR-bin 32 (a high quality MAG), which is annotated to lineage marker set UID3971 by CheckM (selective for Accumulibacter, Dechloromonas and Azospira, all Rhodocyclaceae). See Figure 7b. Examination of the alignments between LR-chromosome 1 and the closely related bin.31 (*sstat* = 0.76) show that its SR-contigs fill a major gap in the coverage of LR-chromosome 1 that is not covered by the members of bin.32 (Figure 7c), suggesting that bin.32 and bin.31 should be a single bin. The closest reference/draft genome identified by MEGAN-LR, GCA 000584975.1 (Candidatus Accumulibacter sp. SK-02), gives an *sstat* of 0.90).

LR-chromosome 3 is contained within the MEGAN-LR taxonomic bin annotated to Chlamydiia, and is covered by contigs from SR-bin 35 (*sstat* = 0.98). The latter bin is a high quality MAG, annotated by CheckM to UID2982, which selects for members of phylum Chlamydiae and phylum Verrucomicrobia. We confirmed the that LR-chromosome 6 (and SR-bin 35) are members of phylum Chlamydiae using a Minimap2 analysis against all extant reference or draft genomes in the PVC superphylum (data not shown). See Figure 7d.

LR-chromosome 4 is contained within the MEGAN-LR taxonomic bin annotated to Gammaproteobacteria and is covered by contigs (*sstat* = 0.97) from SR-bin 43 (high quality MAG), which is itself annotated to Gammaproteobacteria (via UID4266 from CheckM). See Figure 7e.

LR-chromosome 5 is contained within the MEGAN-LR taxonomic bin annotated to *Bacteroidetes bacterium OLB8*, is aligned (*sstat* = 0.98) by SR-bin 66 (high quality MAG), which is also annotated to Bacteroidetes via UID2591 from CheckM. See Figure 7f.

LR-chromosome 6 is contained within the MEGAN-LR taxonomic bin annotated to Rhodospirillales. This LR-chromosome is tiled (*sstat* = 0.95) by contigs from SR-bin 23 (high quality MAG) and which is annotated to order Rhodospirillales by CheckM. SR-bin 23 contains a full length 16S, which Silva assigns to the genus Defluviicoccus (a member of order Rhodospirillales). See Figure 7g.

LR-chromosome 7 is contained within the MEGAN-LR taxonomic bin annotated to *Chlorobi bacterium OLB5*. While there is a good coverage of this LR-chromosome by SR-contigs, these are not contained in in any SR-bin identified by MetaBat. The closest reference genome, GCA 001567546, has a *sstat* value of only 0.36. See Figure 7h.

## Declarations

Ethics approval and consent to participate

Not applicable.

## Consent for publication

Not applicable.

## Availability of data and material

Data from this paper study are available from the NCBI via BioProject accession PRJNA509764 and Short Read Archive accessions SRX5120474 and SRX5126404 for long and short read data, respectively.

DIAMOND, including all modifications introduced in this paper, is open source and available here:

https://github.com/bbuchfink/diamond.

MEGAN Community Edition, including the algorithm for frame-shift correction introduced in this paper, is open source and available here: http://ab.inf.uni-tuebingen.de/data/software/megan6.

## Funding

This work was supported in part by the Singapore National Research Foundation and Ministry of Education under the Research Centre of Excellence Programme, and by a program grant from the Environment and Water Industry Programme Office (EWI), project number 1301-RIS-59. The computational work was partially performed on resources of the National Supercomputing Centre, Singapore (https://www.nscc.sg). The authors acknowledge support by the High Performance and Cloud Computing Group at the Zentrum für Datenverarbeitung of the University of Tübingen, the state of Baden-Württemberg through bwHPC and the German Research Foundation (DFG) through grant no INST 37/935-1 FUGG and grant no HU 566/12-1.

### Acknowledgements

We thank Daniela Drautz-Moses and colleagues for Illumina library preparation and short read sequencing.

## Author contributions

GQ developed and performed the enrichment reactor experiment and obtained samples IB designed and performed the sequencing experiment. RBHW designed the analysis strategy. KA, CB, SB, AG and RBHW performed data analysis. BB implemented frame-shift alignment and range-culling in DIAMOND. DHH implemented frame-shift correction in MEGAN. DHH and RBHW wrote the manuscript and all other authors contributed.

## Supplementary information

*Table S1* Summary of LR-read data, file Table-S1.txt.

*Table S2* Summary of LR-contig data, file Table-2.txt.

*Table S3* Summary of LR-contig taxonomic bins computed using DIAMOND+MEGAN-LR, in file Table-S3.txt.

*Table S4* CheckM results for all 106 LR-contig taxonomic bins, in file Table-S4.txt.

*Table S5* Potential break-points in seven LR-chromosomes, inferred as locations that have no short read clone coverage, in file Table-S5.txt.

*Table S6* Summary of short read data, in file Table-S6.txt.

*Table S7* Summary of short read assembly binning using MetaBat, file Table-S7.txt.

*Figure S1* Plot of LR-contig length vs sstat; highlighting pairs of LR-chromosomes/contigs and SR bins and/or references that show high levels of alignment, file Figure-S1.pdf.

