## Supplementary figures and images for "Annotated bacterial chromosomes from frame-shift-corrected long read metagenomic data"

### Supplemental Figure S1

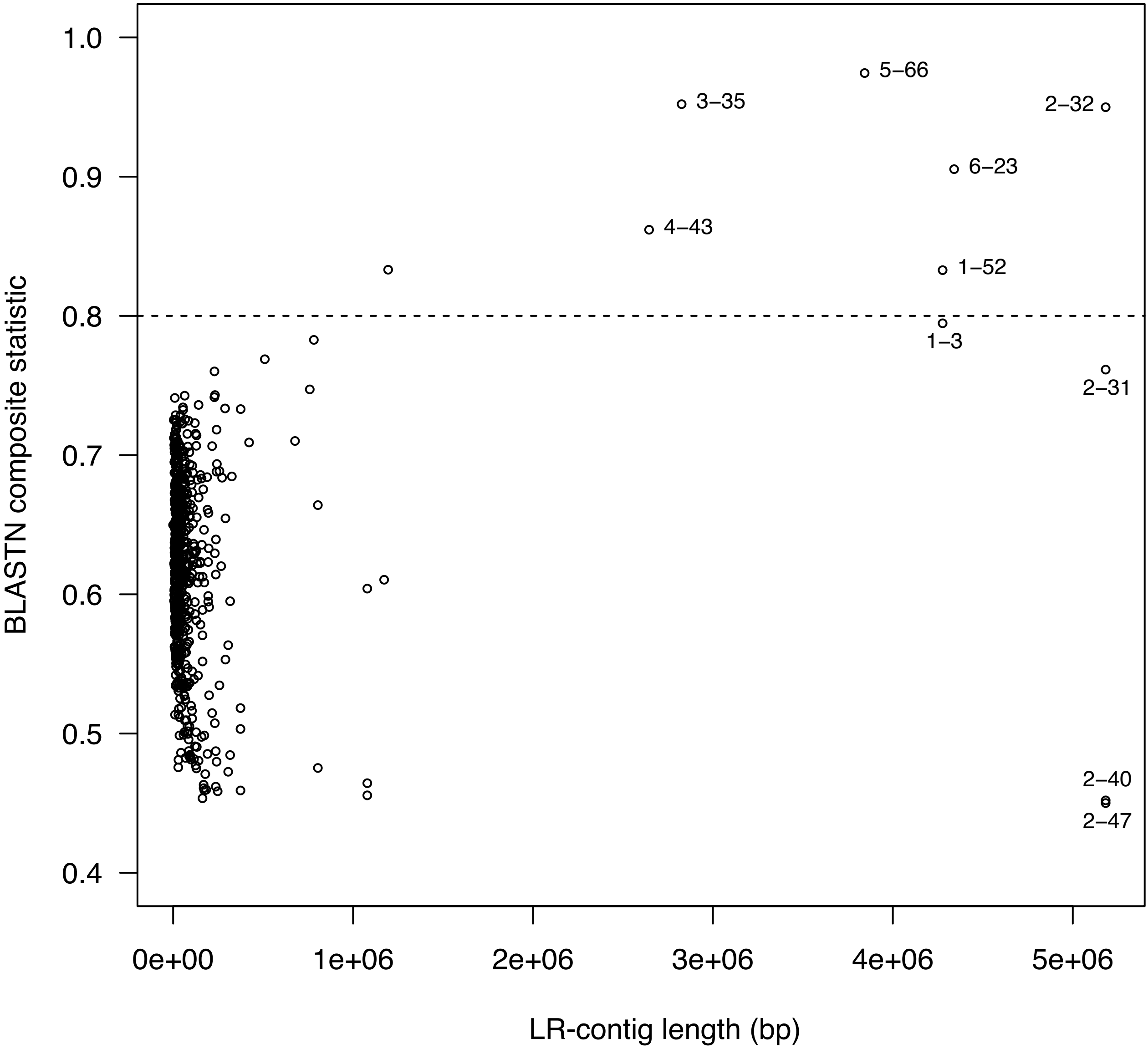
